# Resolving deleterious and near-neutral effects requires different pooled fitness assay designs

**DOI:** 10.1101/2022.08.19.504558

**Authors:** Anurag Limdi, Michael Baym

## Abstract

Pooled sequencing-based fitness assays are a powerful and widely used approach to quantifying fitness of thousands of genetic variants in parallel. Despite the throughput of such assays, they are prone to biases in fitness estimates, and errors in measurements are typically larger for deleterious fitness effects, relative to neutral effects. In practice, designing pooled fitness assays involves tradeoffs between the number of timepoints, the sequencing depth, and other parameters to gain as much information as possible within a feasible experiment. Here, we combined theory, simulations, and reanalysis of an existing experimental dataset to explore how assay parameters impact measurements of near-neutral and deleterious fitness effects. We found that sequencing multiple timepoints at relatively modest depth improved estimates of near-neutral fitness effects, but systematically biased measurements of deleterious effects. We identified a theoretical lower bound for estimates from bulk fitness assays, and showed that increasing sequencing depth, and reducing number of timepoints improved resolution of deleterious fitness effects. Our results highlight a tradeoff between measurement of deleterious and near-neutral effect sizes for a fixed amount of data and suggest that fitness assay design should be tuned for fitness effects that are relevant to the specific biological question.

## Introduction

Accurate fitness measurements are central to questions in experimental evolution, quantitative genetics, and functional genomics. Traditional methods include approaches such as estimating maximum growth rate from growth curves (Hall et al., 2014) and quantifying colony sizes from spot assays (Baryshnikova et al., 2010). An alternative, and increasingly used, approach is a competitive fitness assay in which a reference strain with known fitness is competed directly with a test strain. The relative fitness of the test strain can be inferred from its change in frequency compared to the reference, with either colony counts (Lenski et al., 1991) or fluorescence as a readout (Breslow et al., 2008; Thompson et al., 2006). However, pairwise competition assays are lower throughput and challenging to scale to thousands of measurements.

Competitive fitness assays can be adapted from measuring fitness of a single test strain per assay to measuring fitnesses of several thousand strains in parallel. This typically involves uniquely tagging each strain with a DNA barcode and tracking changes in the frequency of the barcodes over time using deep sequencing (Smith et al., 2009, 2010). Applications of such sequencing based fitness measurements and phenotyping range from CRISPR (Shalem et al., 2014; Wang et al., 2014) and transposon mutagenesis screening for essential genes (van Opijnen and Camilli, 2013; Wetmore et al., 2015), genetic interaction screens (Du et al., 2017; Jaffe et al., 2017), deep mutational scanning of proteins (Fowler and Fields, 2014; Fowler et al., 2010; Stiffler et al., 2015), codon usage (Kelsic et al., 2016), fitness measurements of thousands of adaptive mutations from evolution experiments (Venkataram et al., 2016), genetic crosses (Nguyen Ba et al., 2022) and natural variants (Carrasquilla et al., 2022).

Despite their scalability, highly parallel sequencing-based fitness assays are prone to biases in estimation of fitness. Li et al. demonstrated that fold enrichment based fitness metrics cannot be quantitatively compared across pools of strains with different underlying distributions of fitness effects, and developed FitSeq, a fitness estimation method that accounts for changes in the mean population fitness over time (Li et al., 2018). Their simulations also showed that measuring multiple timepoints makes fitness estimates more robust to changes in the distribution. However, uncertainty in fitness measurements depend on the true fitness and are typically worse for more deleterious fitness effects. Consequently, it is not evident if parameter regimes improving resolution of fitness measurements are the same regardless of the true fitness effect under investigation.

Here, we combined theoretical error bound estimates, simulated fitness assays for a range of experimental regimes, and reanalysis of a deeply sequenced transposon mutagenesis dataset to explore how experimental parameters impact uncertainty in fitness measurements across a wide range of fitness effects. Some of the results presented here have appeared in other work and are cited when appropriate (Li et al., 2018; Robinson et al., 2014); the purpose of this paper is to combine both our findings and these existing insights to derive recommendations for designing pooled sequencing based fitness assays.

### Note about terminology and key assumptions

While we refer to fitness effects of mutations throughout the paper, the results can be extended to any collection of strains, for instance, derived from an evolution experiment, or from natural variation. We define fitness of a mutation with two timepoints, as

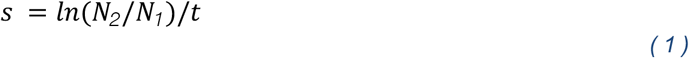

Where N_1_ and N_2_ are the number of read counts before and after selection, and t is the number of generations of selection. This definition can be readily generalized to multiple timepoints as the slope of the linear regression of ln(frequency) vs number of generations of selection in fitness assay. Under this definition, a neutral mutation has a fitness of 0, and an unviable mutation (say loss of an essential gene) has a fitness effect of -ln(2) = -0.693 (Chevin, 2011).

In our simulations, we make some key simplifying assumptions. First, we assume that all mutant trajectories start with the same number of read counts. In experiments, the initial counts are often over-dispersed relative to a Poisson distribution, and the median is less than the mean. Second, we assume that the passage bottleneck size is large enough that genetic drift is minimal. Our rationale for these assumptions is to estimate errors assuming a “best-case” experimental scenario, and to help build intuition for a baseline amount of measurement uncertainty.

## Results

### Errors in fitness measurements depend on the true fitness of a mutant

We first explored how the uncertainty in fitness estimates depends on the “true” fitness of a mutation. We estimated approximate error bounds in fitness estimates assuming that the number of reads at the before and after selection are Poisson distributed. We assumed that there are N_1_ reads before selection and N_2_ reads after selection with N_2_ defined as:

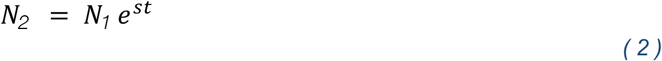

where s is the fitness effect of the mutation, and t is the number of generations of selection. The upper bound is defined as, and the lower bound is defined as:

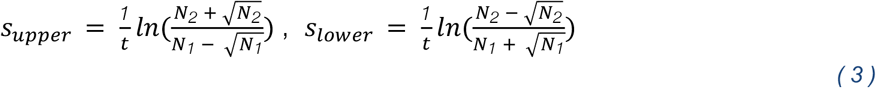

The rationale for these bounds is that for an expected number of reads *N*, the variability due to Poisson noise is 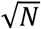. The upper bound in fitness would then be the log ratio of the highest possible number of reads after selection and the least number of reads prior to selection. The lower bound is similarly the opposite. The error in measurement can then be approximated as:

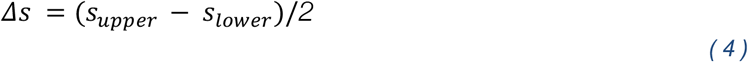

We observe that for more deleterious mutations, the error increases monotonically (Fig 1A). We replicate this observation in simulated fitness assays where we start with an initial number of reads N_1_. We draw from a Poisson distribution with mean N_2_, as defined above. For each mutation we simulate five replicates for a single bottleneck and define the error as the standard error of mean (S.E.M.) of the fitness of the five replicates. Like the error bounds estimated before, we observe that errors are consistently larger for more deleterious mutations (Fig 1B). We turned to a deeply sequenced transposon sequencing dataset of *E. coli* B REL606 from our previous work (Limdi et al., 2022). We found a statistically significant negative correlation between the estimated fitness of disrupting a gene and the error in the estimate (*p*-value < 0.001, Fig 1C); this pattern was more evident when we binned by mutant effect sizes (Fig 1D). This was consistent with the result that FitSeq errors are larger for deleterious mutations (Li et al., 2018). Because errors were dependent on the effect size of the mutation, we decided to explore how experimental parameters impact both near-neutral fitness effects and deleterious fitness effects separately.

**Figure 1:**
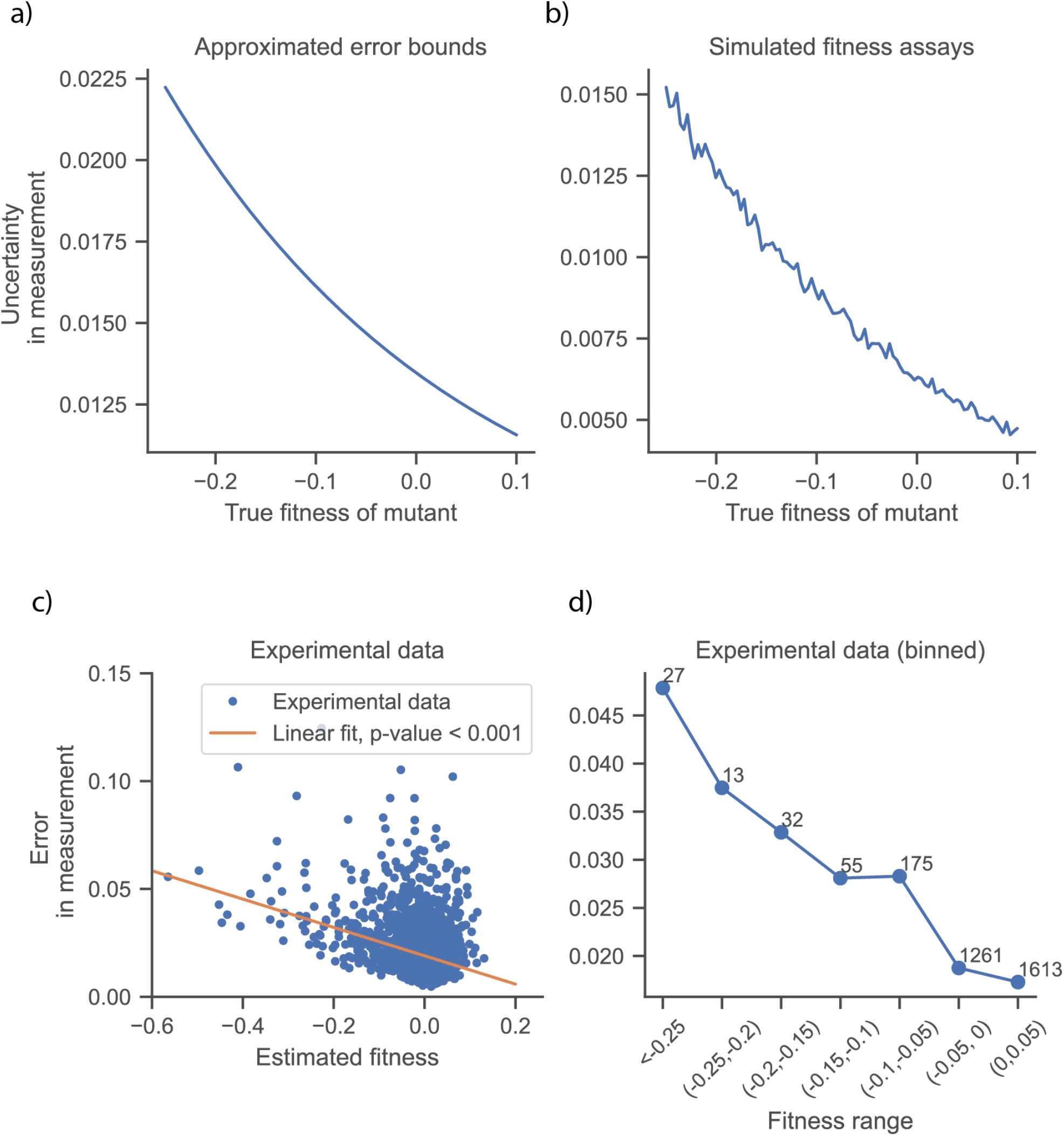
Errors in measurements depend on the true fitness of the mutation. a) Approximate error bounds (equations 3, 4) calculated using Poisson noise as a function of true fitness. Parameters used: number of generations = log_2_(100), sequencing depth per mutation = 500. b) Uncertainty in measurements, defined as the standard error of mean of 5 replicates. Values plotted are the average of fitness assays for 1,000 mutations. Parameters: number of generations = log_2_(100), sequencing depth per replicate = 100. c) Error in fitness measurements (defined as standard error of mean) from a transposon sequencing dataset of *E. coli* B REL606, using two timepoints. d) Same data as in c) binned by fitness effects. Annotations above points indicate the number of genes in the bin.

### Near-neutral mutants require more time for fitness effects to exceed measurement noise

We simulated fitness assays with different sequencing depths, and with varying number of timepoints. More specifically, we counted mutant abundances at multiple timepoints and used all the information to estimate the fitness effect. We observed that as sequencing depth increases, the error decreases, albeit with diminishing returns: increasing depth from 25 to 50 X had a much bigger impact than from 50X to 200X (Fig 2A). This was consistent with lower errors in Bar-seq fitness estimates with increasing sequencing depth (Robinson et al., 2014). Increasing the number of timepoints improved measurements but also with diminishing returns (Fig 2B). We simultaneously varied the sequencing depth, and number of timepoints (Fig 2C) and observed a tradeoff between number of timepoints and sequencing depth; fewer timepoints at lower sequencing depth led to similar measurement errors.

**Figure 2:**
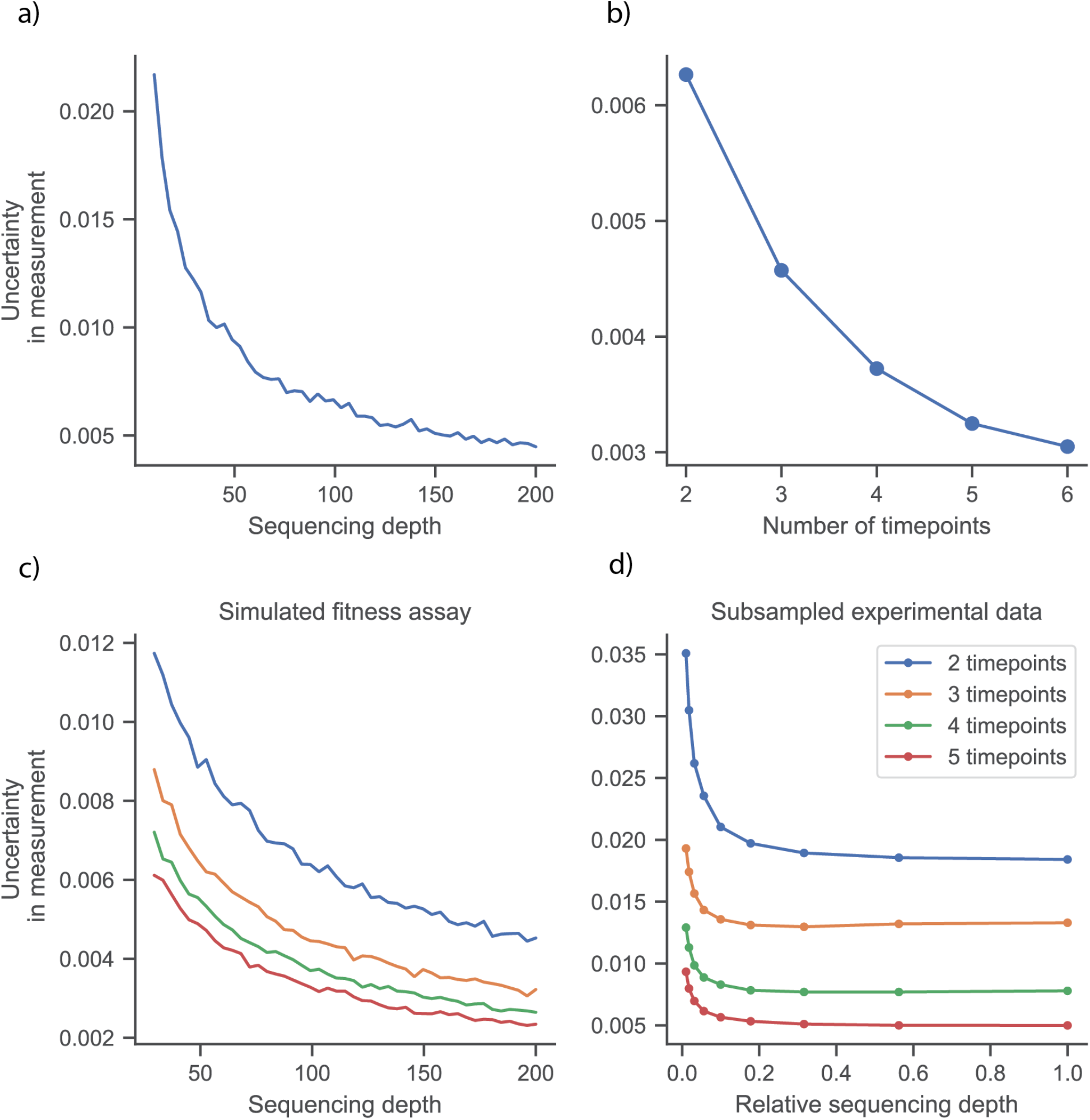
Impact of sequencing depth and number of timepoints on uncertainty in fitness measurements for neutral mutations, with s=0. a) Uncertainty as a function of sequencing depth. Fixed parameters: number of timepoints 2, and number of generations: log_2_(100). b) Uncertainty as a function of timepoints, Fixed parameters: number of timepoints 2, and number of generations: log_2_(100). c) Uncertainty as a function of sequencing depth and number of timepoints, with fixed parameter: number of generations: log_2_(100). In a,b,c) Each point is an average of simulations for 1,000 genes with 5 replicates each. d) Reanalysis of transposon sequencing data with downsampling number of reads (relative sequencing depth) or using a subset of the timepoints for fitness estimation. Since there is no ground truth of neutrality, we averaged over the measurement errors for genes in the range of (−0.05, 0.05).

Next, we explored how tuning the depth and number of timepoints changes errors in the *E. coli* transposon sequencing dataset: we reanalyzed the dataset with down-sampling up to a factor of 10 and using a subset of the timepoints. Like the simulations, we found that the number of timepoints improves the measurement error. However, we observed that the errors depend very weakly on sequencing depth (Fig 2D). This may be due to additional sources of noise (such as bottleneck size, PCR amplification) which set a lower bound on measurement error. Our reanalysis suggested that ∼5-10 times less sequencing depth could have led to similar resolution for near-neutral fitness effects. Further, measurements over five timepoints at 10-fold down-sampling of the sequencing data led to much smaller errors than two timepoints without down-sampling. Note that this corresponds to a fourfold reduction in total number of reads. These results show that for a fixed number of sequencing reads, better resolution of near-neutral fitness effects is obtained with more timepoints and fewer reads per time point.

### For deleterious mutants, error is primarily driven by stochasticity of extinction

Next, we investigated to what extent this intuition also held true for deleterious fitness estimates. For calculating fitness for deleterious mutations from multiple timepoints, we set two constraints: we only calculated fitness effects for replicates that do not die out (i.e., number of counts is above zero throughout). We also required that for each mutation, we were able to calculate fitness for at least two replicates.

We first simulated fitness assays for a range of fitness effects for two timepoints. Errors were consistently larger for deleterious mutations; for the same average error as a neutral mutation, we would need much higher sequencing depth (Fig 3A). Next, we examined how increasing the number of timepoints impacts the measurement error for different fitness effects using the approximate error bounds presented in equation 3 and 4. Consistent with our observations in Figure 2B, errors for near-neutral mutations decrease monotonically with more timepoints. However, for s = -0.2, errors increase after 4 timepoints and for s = -0.3, errors increase monotonically with timepoints (Fig 3B). We confirmed these observations in simulated mutant trajectories for deleterious mutations with slight fitness deficits (s = -0.1, -0.15) for different number of timepoints and sequencing depth. In contrast to neutral mutations, we find that there is no monotonic decrease in error with more timepoints for deleterious mutations; in fact, errors remain unchanged (Fig 3C, compare 3 to 4 timepoints), or increase with more timepoints (Fig 3D). These results suggested that more timepoints can paradoxically make measurements of deleterious mutations more uncertain.

**Figure 3:**
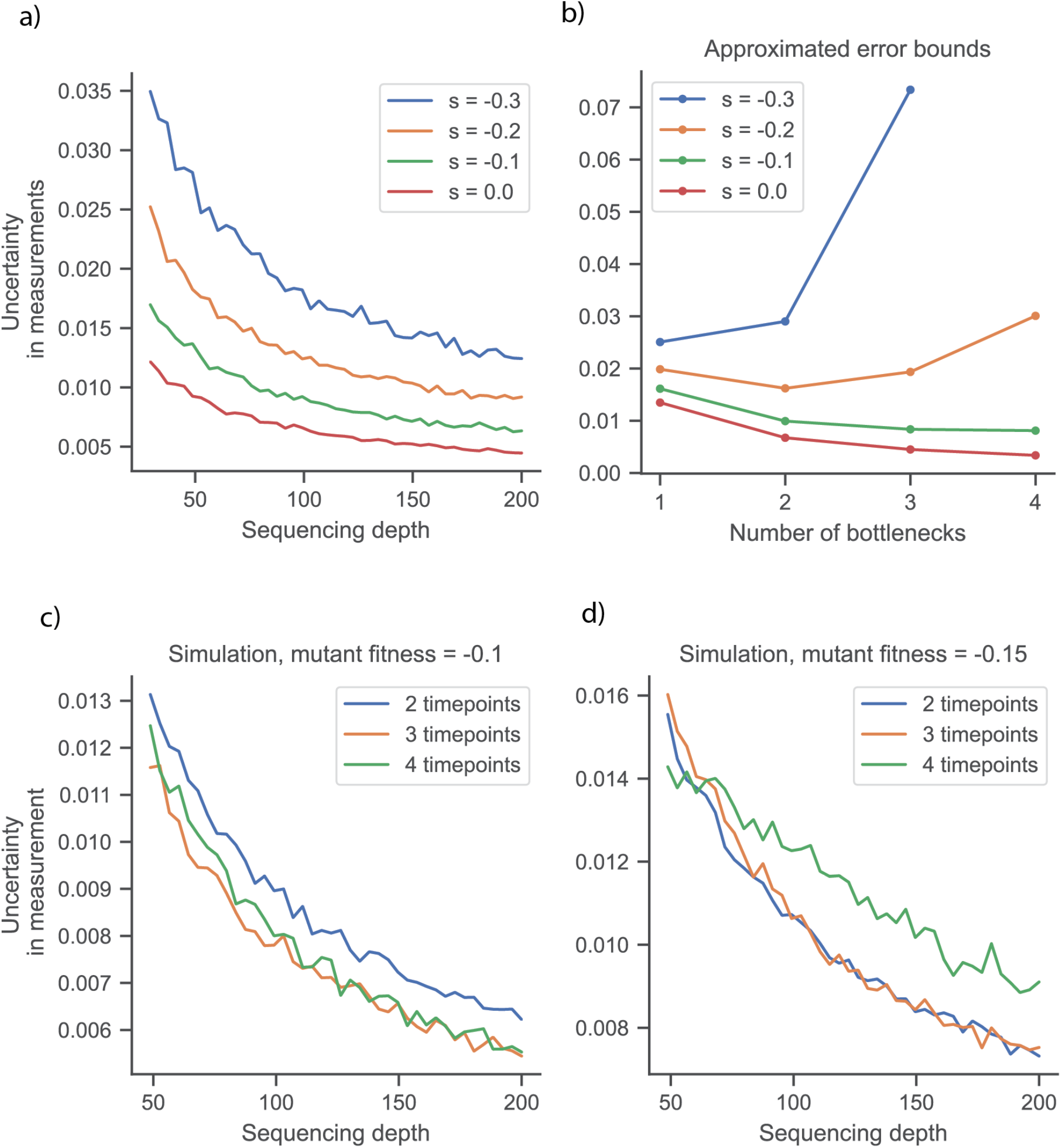
Measurements of deleterious mutations improve with higher depth of sequencing but can worsen with more timepoints. a) Uncertainty as a function of sequencing depth for a range of mutant fitness effects in simulated fitness assays. Fixed parameters: number of timepoints 2, and number of generations: log_2_(100), number of replicates per mutation: 5. b) Approximate measurement error bounds as calculated from equations 3, 4. Fixed parameters: sequencing depth per mutation: 500. c, d) Errors for deleterious fitness effects, calculated using simulated fitness assays for different number of timepoints. Note: in a, c, d) for each parameter combination, we simulated fitness assays for 1,000 mutations (with 5 replicates each) and plotted average error (standard error of mean) from surviving mutant trajectories.

We investigated why error increased with more timepoints. We hypothesized that as the expected counts for deleterious mutations decreased exponentially with time, the number of surviving replicates also decreased, leading to higher uncertainty in the measurement (S.E.M. 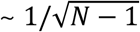). For mutations where we calculated a fitness effect, we counted the number of surviving replicates, as a function of sequencing depth and number of timepoints. Consistent with this hypothesis, we observed that the number of surviving trajectories declined with time (Fig 4A). This trend was particularly evident with stronger deleterious effects.

**Figure 4:**
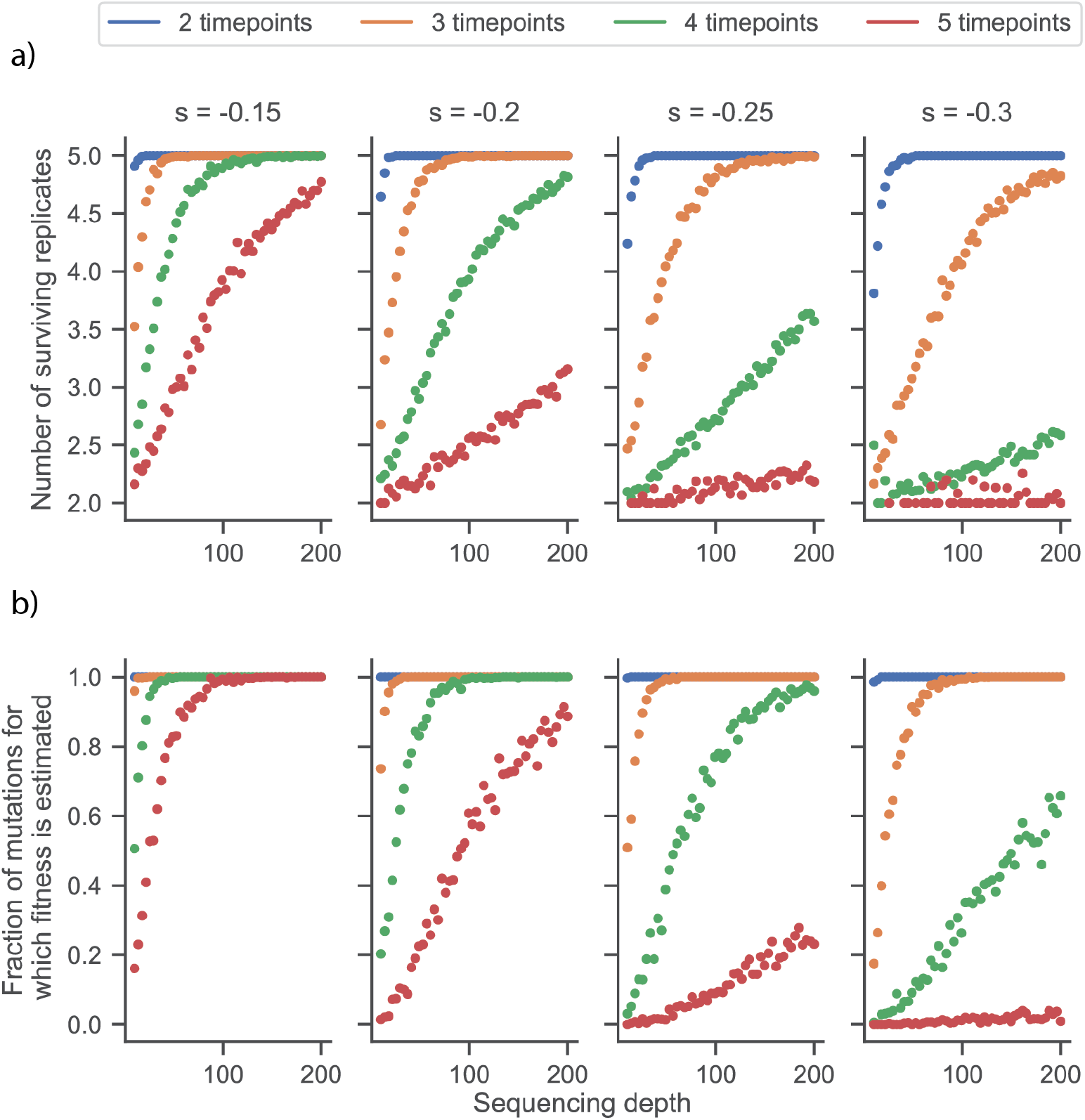
Increasing number of timepoints leads to diminishing information from fitness assays. a) Average number of surviving replicates from simulated fitness assays as a function of sequencing depth and number of timepoints. b) Fraction of mutations with at least two surviving replicates as a function of sequencing depth and number of timepoints. Fixed parameters: number of generations between timepoints: log_2_(100).

Lastly, we explored how the fraction of mutations with at least two surviving replicates (our criterion for calculating fitness) varied with sequencing depth and number of time points (Fig 4B). We observed that for slightly deleterious fitness effects (s = -0.15), the survival probability remained high for five timepoints, and then started declining. In contrast, for more strongly deleterious effects (s=-0.3) was very low after three timepoints, even at relatively high sequencing depths. This suggested that adding more time points yielded progressively less usable information for estimation of deleterious fitness effects. Taken together, these results showed that measurement errors decreased with sequencing depth, but often increased with number of timepoints due to fewer surviving trajectories and therefore usable information from the fitness assay.

### A theoretical lower bound on detectable deleterious fitness effects

While the amount of information that can be used for estimating fitness decreased with time, we wondered whether the remaining surviving trajectories could be used to accurately estimate fitness. On comparing the calculated fitness from these surviving simulated trajectories with the true fitness, we found that fitness was consistently under-estimated for 4 and 5 timepoints. Moreover, this estimate did not change appreciably with greater sequencing depth (Fig 5 A, B).

**Figure 5:**
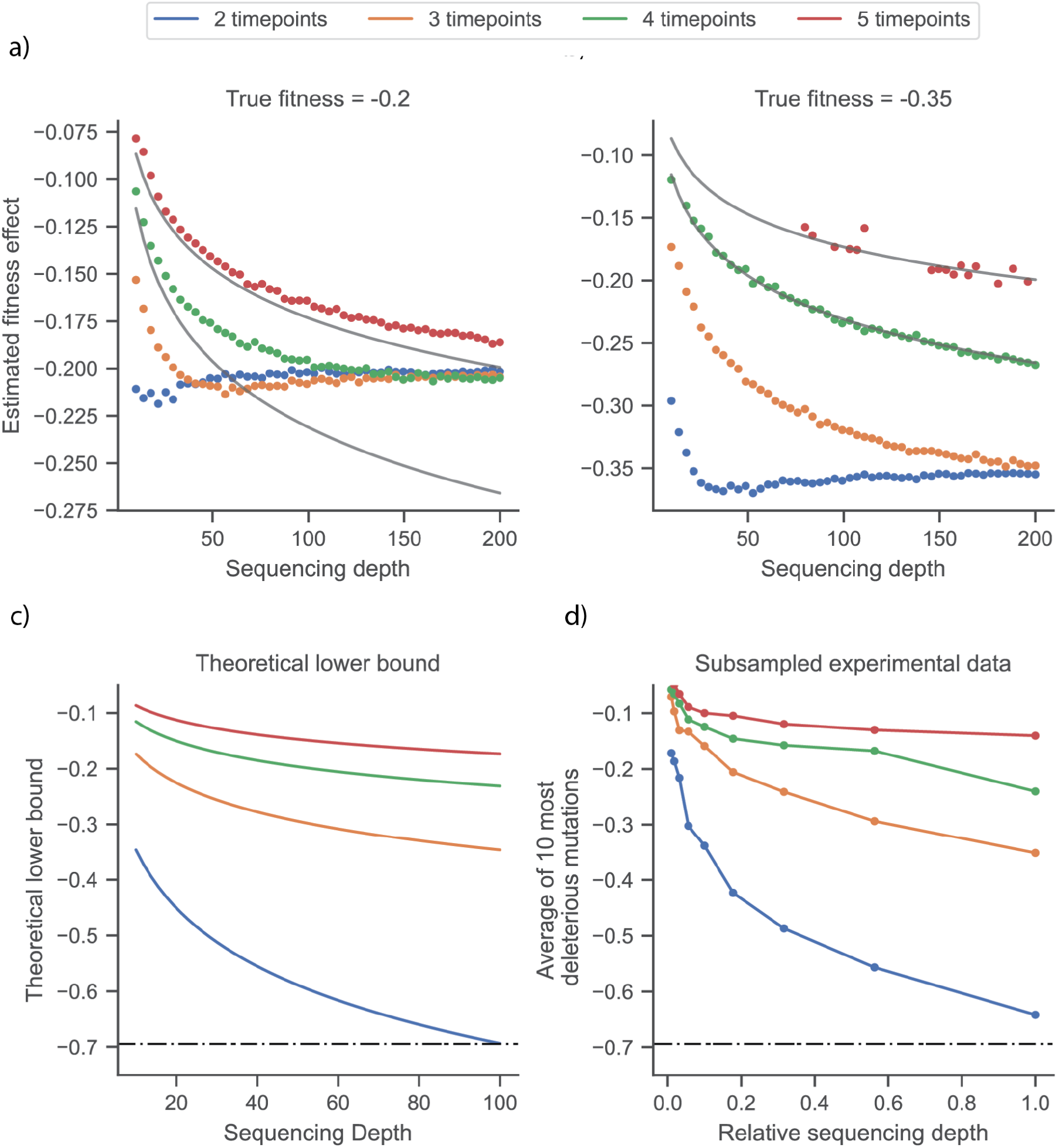
A theoretical lower bound on estimates from pooled fitness assays. a,b) We simulated 1,000 mutant trajectories (5 replicates per mutant) and estimated fitness trajectories met the constraints for estimating fitness effects: at least two replicates per mutation survive; i.e. number of counts does not go to zero. The gray lines indicate theoretical error bounds for 4 and 5 timepoints. c) Plot of theoretical error lower bounds (equation 5) as a function of sequencing depth and number of timepoints. d) Average of the 10 most deleterious fitness effects estimated for a set of parameters (downsampling and timepoints). We use this as a proxy for the resolution of deleterious effects in bulk fitness assays. Dotted line in c,d) indicates the fitness of an unviable mutation, -ln(2).

This occurred because we calculated fitness for surviving trajectories only, which represented a biased subset of the available data. If over half the trajectories do not survive, then it would be quite likely that the number of counts for the surviving trajectories is 1 (assuming Poisson distributed counts). Then, the most deleterious fitness effect obtained from the data would be:

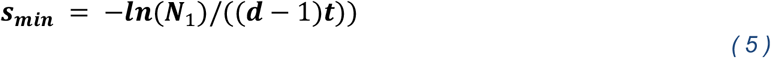

Here, N_1_ is the number of initial counts, and t is the number of generations of selection between timepoints interval, and d is the number of timepoints. When we plotted this against the simulated data, we saw a strong concordance for fitness effects below the theoretical bound, particularly for highly deleterious effects, more timepoints, or lower depth of sequencing, where the counts at the end of the assay are usually 0 or 1 (Fig 5 A, B). Any true fitness below this lower bound was either not estimated at all if every mutant trajectory disappeared to 0 (note the sparsity of points for five timepoints, in red, for a true fitness of -0.35, Fig 5B), or when estimated, converged to this lower bound.

To verify this observation in experiments, we calculated the average of the ten most deleterious fitness effects in the reanalyzed transposon sequencing data, as a proxy for the resolution of deleterious fitness effects and compared against the theoretical bound for different numbers of timepoints (Fig 5C). We required that a trajectory must survive to be included in fitness calculations. We find that the most deleterious effect detected depends strongly on the number of timepoints, and on sequencing depth: a 5-fold decrease in depth has a much larger effect on resolution of deleterious effects than it impacts errors for near neutral mutations (Fig 5D).

## Discussion

We found that sequencing more time points at relatively lower sequencing depth, as opposed to fewer timepoints at very high depth, improves resolution of near-neutral fitness estimates. Conversely, for deleterious fitness effects, with more time points there is less new, usable information obtained, as these variants are depleted over time. In fact, estimates obtained from multiple timepoints can be systematically biased as resolution of deleterious effects is limited by a theoretical lower bound on fitness that can be estimated using pooled assays. We find that resolution of deleterious effects improves with increased sequencing depth and fewer generations of selection, in contrast to measurements of near-neutral mutations. Our results highlight that the timescale of sampling in fitness assays should be tuned to the timescale of change in mutant frequencies. Moreover, they suggest that there is no combination of experimental parameters that optimally resolves both ranges of fitness effects for a fixed amount of data.

A limitation of our simulations is that we make several simplifying assumptions in modeling fitness. We do not consider amplification noise during PCR, bottleneck noise, and do not account for heavy tailed distribution of initial read counts. These factors likely contribute to higher measurement noise. For a more detailed analysis of this concern, we recommend (Li et al., 2018). Further, we do not aim to prescriptively specify experimental parameters or error bounds but discuss factors influencing uncertainty in fitness measurements and explore the tradeoffs involved as a starting point for experimentalists to tune fitness assay design.

Our simulations and reanalysis of transposon sequencing data, combined with previously published results, can be distilled into principles for experimental design:

1. *Identify fitness effects that are relevant for the biological question at hand* Errors in measurements depend on the true fitness of the mutations, and parameters that improve near-neutral fitness measurements are sub-optimal for assaying deleterious fitness effects.
2. *For measurements near neutrality, sequence mutant pools at multiple timepoints, with relatively modest sequencing depth* Our reanalysis of a deeply sequenced transposons sequencing data show sequencing more time points at 1/10th the depth leads to better fitness estimates than a single time point. Additionally, the mean fitness of the population can change over time depending on the underlying distribution of fitness effects (Li, et al.). This can lead to biased fitness estimates; for instance, neutral mutations may appear deleterious without any correction. Quantifying mutant abundance over multiple timepoints allows for use of methods such as FitSeq to correct for this bias.
3. *For measurements of deleterious mutations, sequence at high depth, for fewer timepoints and fewer generations* Including multiple timepoints does not add meaningful information as deleterious mutations will disappear over time. Further, including data from multiple timepoints can lead to systematically biased estimates if the expected number of counts is close to 0 at any time point. As a starting point for parameters, we recommend using sequencing depth and number of timepoints such fitness effects of interest are above this theoretical bound. Lastly, we recommend (from experience) always sequencing the mutant pools prior to any fitness assay, as deleterious mutations (or variants) disappear from the pool rapidly in a few generations.
4. *Using pilot experiments and simulations to guide fitness assay design* We present an approach for tuning fitness assay design; we suggest performing a pilot fitness assay and sequencing experiment, using simulations as a starting point for experimental parameters, and reanalyzing the data with subsampling (either fewer reads or fewer timepoints). If the errors or resolution of fitness effects of interest do not change with subsampling, it is possible to collect data for more strains/genetic backgrounds with the same total sequencing.

## Statements and Declarations

### Data Availability

Raw sequencing reads have been deposited in the NCBI BioProject database under accession number PRJNA814281. Processed data are deposited on Zenodo (https://doi.org/10.5281/zenodo.6547536), and source code for sequencing pipeline, downstream analyses, and figure generation are available at GitHub (https://github.com/baymlab/2022_Limdi_limits-pooled-fitness-assays)

## Acknowledgments

We thank Fernando Rossine, Eleanor Rand, and Indra Gonzalez Ojeda for feedback and discussion on analysis and figures. A.L. acknowledges support from the Molecules, Cells, and Organisms Graduate Program, Harvard University. M.B. acknowledges support from the NIGMS of the National Institutes of Health (R35GM133700), the David and Lucile Packard Foundation, the Pew Charitable Trusts, and the Alfred P. Sloan Foundation.

